# Salivary oxytocin co-varies with parturition and nursing behavior in domestic pigs (*Sus scrofa domesticus*)

**DOI:** 10.1101/2021.06.11.448025

**Authors:** Liza R. Moscovice, Birgit Sobczak, Ulrike Gimsa, Ellen Kanitz

## Abstract

Domestic pigs produce twice as many offspring as wild boars, but little is known about the effects of selection pressures for increased productivity on pig behavior and welfare. From an evolutionary perspective, producing larger litters is expected to increase parent-offspring conflict, which may help to explain why piglets from larger litters are more likely to die by starvation or infant-crushing. Our goals were to identify behavioral and physiological correlates of variation in maternal care that may be useful as selection criteria in breeding programs to improve pig welfare. We observed eleven sows regularly during their lactational period and recorded their hourly postural changes and the proportion of time that piglets had contact with teats (nursing contact) or any physical contact with the sow (social contact). As a potential physiological indicator of maternal care, we measured oxytocin concentrations in 119 saliva samples from sows following observations. Samples were extracted and measured in an Enzyme Immunoassay. As a biological validation, we measured changes in salivary oxytocin during parturition in an additional eleven sows. Laboratory validations confirmed that the assay is suitable for measuring salivary oxytocin in pigs, and oxytocin increased almost three-fold during parturition. Sows’ nursing contact was positively correlated with social contact and negatively correlated with postural changes. Sow oxytocin concentrations were predicted by their nursing contact, but not by their overall social contact with piglets. We compare our results with current evidence regarding best practice methods for salivary oxytocin measurements and discuss the potential to use indicators of maternal care as selection criteria to improve pig welfare.

## Introduction

During their domestication, the descendants of wild boars have undergone strong selection pressures for greater productivity [1,2], resulting in a doubling of average litter size at birth, from 5.4 (± 1.5) live born offspring in wild boars [3] to 11.7 (± 3.2) in domestic breeds [4], with the largest litters often containing more than 16 piglets [5]. Less attention has been given to the potential impacts of these selection pressures on the welfare of sows and their offspring [2,5]. From an evolutionary perspective, producing larger litters is expected to exacerbate short-term fitness tradeoffs for mothers, between investing in more versus healthier offspring within a current litter, as well as long-term fitness trade-offs, between investing in current versus future litters [6]. There is evidence that domestic pigs (*Sus scrofa domesticus*) still experience such trade-offs, since high milk production can exceed a sow’s capacity for feed intake, resulting in worsening body condition and increased lesions during lactation, compared to less productive sows [1,6].

Increased parent-offspring conflict over allocation of resources may result in reductions in maternal investment, and in extreme cases infanticidal behavior [7]. This socio-biological perspective can help to explain the evidence that piglets from larger litters are more likely to die by starvation or infant-crushing [8], which are the two greatest risks to piglet survival, together accounting for up to 60% of pre-weaning mortality [9]. These two risks may be inter-related, since piglets who are not receiving milk may be too weak to avoid being overlain by mothers [5], and mothers may selectively direct crushing behavior at offspring who are not receiving milk [8]. High piglet mortalities represent not only a major economic loss but also an ethical problem, which may be addressed by balancing genetic selection for production with selection for additional traits that predict enhanced maternal care.

Even under the highly standardized conditions in conventional pig breeding facilities, domestic sows exhibit inter-individual variation in several behaviors that may influence maternal care, including general activity levels and frequencies of postural changes (e.g. from standing to laying down [10], rates of affiliative social contact with offspring [11] and the frequency and duration of nursing events [12,13]). Nursing behavior is of particular interest for understanding how sows may respond to parent-offspring conflict, since sows lack teat cisternae to store milk. As a result, piglets are only able to feed during brief milk let-down events, which are controlled by the sows. The initiation of milk let-down is typically signaled by sows using rapid ‘nursing grunts’, which attract piglets to the teats [14]. Piglets then massage the teats which helps to trigger milk ejection [15]. After milk ejection, piglets often resume teat massage for several minutes, which has been hypothesized to prime future milk release for piglets [15], who typically show stable teat preferences [16]. Sow nursing frequencies can vary greatly during the first few days post-partum, but subsequently take on a more regular rhythm, with inter-nursing intervals ranging from 30-70 minutes [13,15,17]. Since the udder can only be refilled after milk-letdown has occurred, decreasing the inter-nursing interval is the main mechanism by which milk production can be increased [6,12]. This has been demonstrated experimentally by showing that inducing nursing at shorter intervals was associated with greater milk consumption and greater weight gain in piglets [12].

Characterizing physiological correlates of maternal behavior can help to identify proximate causes of inter-individual variation in maternal investment that may also be amenable to genetic selection. The neuropeptide hormone oxytocin (OXT) is an obvious candidate as a physiological indicator of maternal care. Oxytocin is synthesized in paraventricular and supraoptic nuclei of the hypothalamus and can be released from hypothalamic nuclei to act as a neurotransmitter in the brain, or it can be transported to the posterior pituitary and released peripherally into the blood stream, where it acts as a hormone [18]. Oxytocin has a highly conserved role in mediating various aspects of female reproductive and maternal behavior in mammals [19], including domestic pigs [15,20]. In sows, basal plasma OXT concentrations are negatively associated with the duration of parturition [17,21], and positively associated with piglet growth [22], and udder stimulation is associated with spikes in plasma OXT [20,22,23].

In addition to reflecting physiological processes related primarily to female reproductive and maternal functions, there is increasing evidence that central activation of the oxytocinergic system is sometimes associated with peripheral OXT release [24,25]. There has been a corresponding increase in studies reporting co-variation between peripheral OXT measured in plasma, urine or saliva and salient biological stimuli [26,27]. In human mothers, OXT concentrations in plasma and saliva are correlated [28,29], and both measures positively predict maternal social engagement with offspring [28,30]. In sows, plasma OXT has been linked to maternal body condition and offspring growth, but not to maternal behaviors or mother-offspring interactions [22]. This may be due to the difficulty in obtaining repeated plasma samples from sows, which is often achieved via catheterization, thus requiring specialized personnel, and having a greater potential to cause stress or discomfort to sows.

The goals for this study were to determine whether various measures of maternal behavior were stable and correlated within individuals, varied between sows, and were related to variation in peripheral OXT concentrations. We chose to measure OXT in saliva, since this medium is most suited to repeated sampling of individuals while having the least interference with on-going behavior. Despite initial doubts about the origins and measurement of OXT in saliva [31], subsequent studies have found increases in salivary OXT in humans and other species related to biological events that are mediated by the oxytocinergic system [29,32,33]. Although we now understand more about the various mechanisms by which small molecules like OXT (molecular weight = 1007 Da) may transfer from the blood stream into oral fluid, including via ultrafiltration [34], valid criticisms remain about salivary OXT measurements, including a lack of standardized methods for processing of saliva prior to measurement [35] and a lack of thorough validations when applying the method to new species. Such validations are especially important when measuring salivary OXT using commercially available OXT immunoassay kits, which were initially designed and validated for plasma. Salivary oxytocin measurements have recently been validated in pigs [36,37], but using different methods, and with contradictory results regarding the importance of extraction prior to measurement. An additional goal for our research was to perform a series of analytical and biological validations of our salivary OXT measurements, to help advance the discussion over best practice methods for measuring peripheral OXT in saliva [35,38].

## Methods

### Subjects and husbandry

Subjects were primarily German Landrace sows (n = 17), along with a small number of Angeln Saddleback sows (n = 5). All subjects were born and raised in the experimental pig facility (EAS) of the Leibniz Institute for Farm Animal Biology, Dummerstorf, Germany. Husbandry conditions in the EAS are consistent with those in conventional pig breeding facilities in Germany, as required by the Animal Welfare Ordinance on the Keeping of Farm Animals [39]. No alterations were made for this study. Farrowing is synchronized among several sows, who are housed individually in conventional loose farrowing pens (6 m^2^) beginning on the 105^th^ day of pregnancy. Farrowing pens contain a wood chew toy for enrichment and a heated, covered lying area (90 × 90 cm) for piglets. Rooms have an automatic ventilation system, controlled lighting (12/12 h light/dark cycle, lights on at 0600 h) and are kept at a nearly constant temperature (28 ± 1 °C). While in farrowing pens, sows are fed twice daily with a pregnancy feed (Sow Structure 7.0, Trede & von Pein GmbH, Itzehoe, Germany) until six days prior to birth, at which point they are switched to a lactation feed (Sow Provital, Trede & von Pein GmbH). Water is provided *ad libitum*. From three days prior to parturition until ten days post-birth, a guard rail is employed as a preventative measure to reduce piglet crushing [40], which limits the sows’ available area to approximately 200 x 90 cm. From ten days post-birth, the guard rail is removed and sows have access to the entire farrowing pen, with the exception of the pig laying area, which remains out of reach to the sow. Piglets can move freely in the farrowing pen throughout the lactational period. Piglets have unrestricted access to water and are offered dry food (HAKRA-Immuno-G; Una Hakra, Hamburg, Germany) in addition to milk beginning from 14 days of age. Farrowing pens are cleaned daily, and piglets are weighed weekly by EAS staff. Otherwise piglets and sows are left undisturbed. Piglets are not subjected to tail docking, teeth clipping or male castration. All research procedures were approved under the German Animal Welfare Act (German Animal Protection Law, §8 TierSchG) by the Committee for Animal Use and Care of the Agricultural Department of Mecklenburg-Western Pomerania, Germany (permit LALLF 7221.3-1-003/18).

For the biological validation of OXT, subjects were eleven primiparous sows (nine German Landrace and two Angeln Saddlebacks). Based on decisions by the EAS staff, six of these sows received synthetic prostaglandin injections (PGF Veyx^®^, Provet AG, Lyssach, Switzerland) and three of these same sows also received synthetic OXT injections (LongActon^®^, Vetoquinol AG, Bern, Switzerland), to induce parturition. The other sows remained untreated. For the research on maternal care, we collected behavioral data and saliva samples from an additional eleven primiparous sows (eight German Landrace and three Angeln Saddlebacks) during their first lactational period. We used different subjects from those involved in the biological validation. We collected behavioral data and corresponding saliva samples from these sows 3-4 times weekly from post-birth day 1 until day 27, one day prior to weaning. Based on decisions by the EAS staff, 13.5% (n = 21) of live-born piglets from seven sows were fostered to different mothers within one day of birth, as a method to balance out litter sizes and to optimize birth weights within litters. This practice is widely applied in conventional breeding facilities [41], and both domestic sows and wild boars show willingness to adopt piglets that are introduced shortly after birth [42], a behavior that likely has indirect fitness benefits in natural settings, in which alloparenting may occur among several related females [42].

### Behavioral observations

We used scan sampling methods to record behaviors from several sows within the same time period. Between 4-5 sows were observed for one hour beginning either at 8:30 or 10:30, with the time of observation counter-balanced across days. During each observation period, one of two authors (L.R.M. or B.S.) observed every sow at 2-minute intervals, resulting in 30 scans per day per sow. At each scan interval, an observer moved between each pen in sequence and noted for each sow, their posture (standing, sitting, laying on side with some teats exposed, or laying on belly with no teats exposed), the number of piglets who were in physical contact with teats, and the number of piglets who were in physical contact with any other body part of the sow. We used these data to calculate for each sow her hourly: 1) number of postural changes, defined as the number of shifts from one posture to another across scans; 2) nursing contact, defined as the proportion of scans in which one or more piglets had contact with teats and 3) social contact, defined as the proportion of scans during which one or more piglets had any type of physical contact with the sow (with teats or with other body parts). For nursing and social contact, we did not correct for differences in the number of piglets in contact at each scan, since we expected peripheral OXT to be related to the frequency of affiliative and nursing-related social contact, regardless of the number of piglets involved. We also calculated for each sow their 4) number of nursing events. Following Valros and colleagues [13], a nursing event occurred on scans when more than 50% of the piglets were suckling and ended on scans when more than 50% of the piglets were not in physical contact with the teats anymore. Independent nursing events were defined when there were at least two intervening scans without any teat contact. We did not attempt to distinguish between non-nutritive and nutritive nursing contact, as both were relevant for our research questions. However, other studies have found that only a small percentage of nursing events occurred without milk ejection (e.g. 4%, [12]).

### Salivary oxytocin sampling and measurement

We used SalivaBio^®^ Childrens Swabs (Salimetrics, California, USA), taped to wooden dowels, to collect saliva samples from sows. For the biological validation, samples were collected one to two days prior to parturition and during parturition. Pre-parturition samples were collected in the morning between 9:00–11:00. We attempted to collect the parturition samples between 2–4 hours after the first piglet expulsion, based on other studies indicating peaks in plasma OXT concentrations during this period [17]. For the study on maternal behavior, samples were collected between 9:30–11:45, within fifteen minutes following the end of each hour-long observation period. Due to the delay between exposure to salient stimuli and changes in salivary OXT [32], we expected that the OXT samples would relate to the time period during which the behavioral observations had occurred.

All saliva sampling occurred at least an hour after the morning feeding (at approx. 8:00), and prior to the afternoon feeding (at approx. 14:00), which ensured that sows did not eat prior to sampling. We also ensured that sows did not drink water for 10 minutes prior to sampling. To obtain samples, we entered the pen and placed the swab in the sow’s mouth, attempting to sample between the gum and cheek, or under the tongue, for up to 2 minutes. Saturated swabs were inspected and discarded if there were any signs of blood on the swabs, which occurred in two samples (1.7%). Swabs were placed in polypropylene tubes on ice and centrifuged within 15 minutes of collection at 4° C (15 min, 2,000 x g). The eluent was stored at - 80° C until hormone analyses. We stored multiple aliquots of each sample whenever possible, to use for additional assay validations.

Prior to measurement, we performed a solid phase extraction (SPE) following a protocol modified from Lürzel and colleagues [36]. Samples were thawed on ice and centrifuged at 4° C (5 min, 2,000 x g). We mixed 250 μl of saliva with equal parts 0.1% trifluoroacetic acid (TFA)/water, vortexed, and then centrifuged at 4° C (15 min, 17,000 x g). During preliminary tests, we found that this second centrifugation step in TFA helped to separate out additional particles that may have interfered with OXT measurement. We loaded the supernatant on 30 mg 1 mL HLB cartridges (Oasis Prime^®^, cat no. 186008055, Waters Corporation, MA, USA), primed with 1 mL 99% acetonitrile (ACN), followed by 3 mL 0.1% TFA/water. Samples were washed with 3 mL 0.1% TFA/water and eluted with 3 mL 60% ACN/40% 0.1% TFA. Eluents were evaporated using a Vacuum concentrator (SpeedVac^®^ SC210, Thermo Fisher Scientific, MA, USA), and frozen at −80° C until measurement. We measured OXT using a commercially available Enzyme Immuno-Assay (EIA; Caymen Chemical, kit no. Cay500440, MI, USA). The standard curve ranges from 5.9–750 pg/mL and assay sensitivity is 20 pg/mL. Prior to measurement, we reconstituted samples in 250 μl assay buffer, and then followed the kit instructions. We performed validations of parallelism, accuracy/recovery and reproducibility to confirm that the assay measures the majority of OXT present and is suitable for measuring OXT in pig saliva (see results and S1 Fig). For reproducibility, we measured intra- and inter-assay coefficients of variation in a pooled saliva sample from ten different sows that was stored in multiple aliquots at −80° C and extracted along with test samples.

To assess the impact of extractions on sample measurement, we compared a subset of samples that were collected in duplicate, either extracted or left unextracted and then measured on the same plate. Unextracted samples were thawed and centrifuged. The supernatant was combined with equal parts assay buffer and plated following the kit instructions. Additionally, we compared kit efficacy of EIAs for measurement of salivary OXT in pigs by extracting a subset of samples collected in duplicate and measuring them with both the Caymen assay and the DetectX^®^ Oxytocin EIA from Arbor Assays (kit no. K048-H1, MI, USA).

### Statistical analyses

Statistical analyses were performed using the R statistical software [43]. To assess the relationship between concentrations of duplicate samples analyzed with different assays, or with and without extraction, we ran Spearman correlations (r_s_), using the ‘pspearman’ package [44]. To compare salivary OXT concentrations in sows before and during parturition, we ran Wilcoxon signed-ranks tests, using the package ‘coin’ [45]. To determine whether daily measures of different maternal behaviors were correlated with each other, we computed repeated measures correlations (r_rm_, [46]) using the package ‘rmcorr’ [47]. To determine if sows showed stability in their maternal behaviors during lactation, we used Spearman correlations to compare each sow’s average behavioral measures across three post-birth time periods: days 1–10, 11–19 and 20–27. When performing multiple correlation tests, we used the Bonferroni correction to control for inflated type I error rates.

We built linear mixed models (LMMs) using the package ‘lme4’ [48] to test whether salivary OXT in sows was predicted by their proportion of scans spent in nursing contact, or in any social contact with piglets. Due to a lack of independence between these two behavioral measures, we tested each predictor in a separate model. As control predictors in both models, we included the time of sample collection, the time delay between the last nursing event and sample collection and the number of days since birth. We square-root transformed OXT concentrations, nursing contact and time delay between nursing and sampling to approximate normal distributions, and we z-transformed all predictors, to facilitate interpretability of the results. We included random intercepts for subject and date, and random slopes for nursing contact or social contact within subject and date, to keep the type I error rate at a nominal 5% [49]. We confirmed that models were stable by comparing the model estimates derived from the full data set with those obtained from a model with each subject excluded one by one. We also tested for variance inflation using the ‘vif’ function in the package ‘car’ [50] and found no evidence for collinearity of the predictors (VIFs < 1.16). We used likelihood ratio tests to compare each model to a reduced model excluding the test predictor, and present results of models that differed significantly from the reduced model. Unless otherwise noted, results are presented as mean (± SD) and a *P*-value <0.05 was considered significant.

## Results

### Analytical validations and comparison of assays

For the Caymen assay, tests of parallelism revealed no differences in the slopes between the standard curve and a serial dilution of pooled pig saliva (t = 0.17, p = 0.86, n = 4, S1a Fig). Recovery of OXT in three different pooled samples of known concentrations, each spiked with OXT standards at three different concentrations spanning the linear range (750 pg/mL, 375 pg/mL, 187.5 pg/mL) was 96.4 ± 6.8% (n = 9 spiked samples). The intra-assay CV was 6.7% (n = 11 samples), and the inter-assay CV was 24.5% (n = 6 samples). Although the higher inter-assay CV implies some plate-to-plate variation in measurements, this should have little effect on our results, since we balanced samples from different subjects across plates. Extracted and unextracted samples measured on the same Caymen plate were not correlated (r_s_ = 0.36, n = 10, p = 0.31). Oxytocin concentrations in duplicate samples extracted using the same method and measured with both Caymen and Arbor assays were moderately correlated (r_s_ = 0.44, n = 36, p = 0.045). However, a comparison of the validations between the two assays indicated lower parallelism (S1b Fig) and lower reproducibility (intra-assay CV = 26%, n = 12 samples) in the Arbor assay in comparison with the Caymen assay.

### Oxytocin during parturition

Salivary OXT concentrations increased almost three-fold during parturition, compared with one to two days before (553.2 ± 372.1 pg/mL vs. 198.6 ± 102.5 pg/mL, Wilcoxon, N = 11, Z = −2.41, p = 0.02, Fig 1). Although parturition was associated with increases in salivary OXT concentrations in 91% (n = 10) of sows, relative changes in OXT varied greatly among individuals (Table 1).

**Fig 1.**
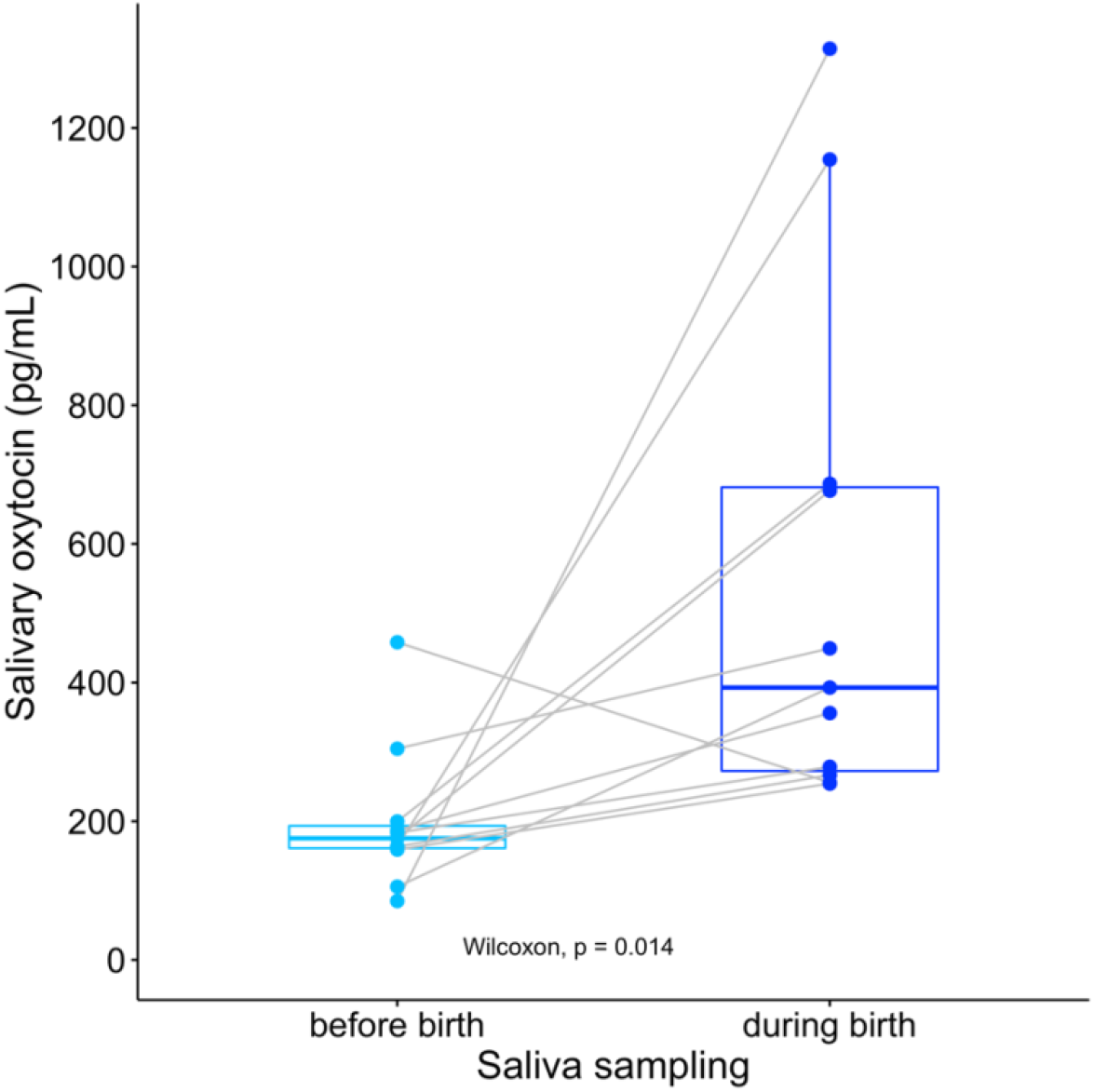
Changes in salivary oxytocin concentrations during parturition. Oxytocin was measured in eleven sows one to two days prior to parturition and during parturition.

**Table 1.** Relative changes in salivary OXT concentrations during birth. (OXT post-birth – OXT pre-birth / OXT pre-birth). DL= German Landrace, AS= Angeln Saddleback, PGF= synthetic prostaglandin, Long= synthetic OXT

| Sow ID | Breed | Parturition date | Treatment during parturition | Relative change in salivary OXT during birth |
| --- | --- | --- | --- | --- |
| 6076 | DL | 20-Jun-19 | none | 14.4 |
| 5989 | DL | 21-Jun-19 | none | 0.63 |
| 5986 | DL | 22-Jun-19 | none | 0.60 |
| K6254 | AS | 14-Sep-19 | none | 0.52 |
| K6255 | AS | 23-Oct-19 | none | - 0.44 |
| 6073 | DL | 3-Sep-20 | PGF | 2.43 |
| 6263 | DL | 3-Sep-20 | PGF + Long | 0.47 |
| 6271 | DL | 3-Sep-20 | PGF + Long | 0.91 |
| 6361 | DL | 3-Sep-20 | PGF | 2.72 |
| 6364 | DL | 3-Sep-20 | PGF | 2.86 |
| 6369 | DL | 3-Sep-20 | PGF + Long | 6.07 |

### Maternal behaviors during lactation

Sows had 5.5 ± 3.2 postural changes per hour and were in social contact with piglets during 55.8 ± 25.2% of hourly scans. Sows exhibited a great degree of variation in their hourly nursing contact with piglets, which ranged from 4–40% of scans (mean= 17.8 ± 17.1% of scans, Table 2). Sows had 1.3 ± 0.7 nursing events per hour (where > 50% of piglets were in nursing contact). Sow hourly nursing contact was positively correlated with their hourly number of nursing events (r_rm_ = 0.34, df = 109, p < 0.001, 95% CI = 0.16– 0.49), and negatively correlated with their hourly number of postural changes (r_rm_ = −0.28, df = 109, p = 0.003, 95% CI = −0.48– −0.10). As expected, nursing contact was correlated with overall social contact (r_rm_ = 0.61, df = 109, p < 0.0001,95% CI = 0.48– 0.72). Hourly proportion of scans in social contact was also positively correlated with hourly number of nursing events (r_rm_ = 0.37, df= 109, p < 0.001,95% CI= 0.20- 0.52). Other behaviors were not correlated (p > 0.22, Table 3). Sows exhibited intra-individual stability in nursing contact when comparing between later observation periods (days 11–19 vs. days 20–27, r_s_ = 0.97, n = 11, p < 0.0001). Sows also exhibited intra-individual stability in social contact, but only at earlier observation periods (days 1–10 vs. 11–19, r_s_ = 0.91, n = 11, p < 0.0001). Number of nursing events and postural changes were not stable across different time periods (S1 Table).

**Table 2.** Variation in behavior and salivary oxytocin concentrations during lactation. Sows are arranged based on their average hourly proportion of scans in nursing contact (predictor in model), from lowest to highest. DL= German Landrace, AS= Angeln Saddleback.

| Sow ID | Breed | Postural changes<br>(Mean ± SD) | Proportion<br>social contact<br>(Mean ± SD) | Proportion<br>nursing contact<br>(Mean ± SD) | Nursing<br>events<br>(Mean ± SD) | OXT pg/mL<br>(Mean ± SD) | No.<br>piglets<br>weaned |
| --- | --- | --- | --- | --- | --- | --- | --- |
| 6404 | DL | 6.27 ± 3.82 | 0.69 ± 0.18 | 0.04 ± 0.03 | 1.00 ± 0.45 | 214.67 ± 70.62 | 13 |
| 6381 | DL | 3.37 ± 2.54 | 0.47 ± 0.23 | 0.05 ± 0.08 | 0.91 ± 0.70 | 218.88 ± 67.92 | 14 |
| 6398 | DL | 4.82 ± 2.14 | 0.79 ± 0.16 | 0.08 ± 0.08 | 1.55 ± 0.52 | 155.75 ± 63.04 | 12 |
| 6284 | AS | 3.82 ± 2.09 | 0.30 ± 0.10 | 0.11 ± 0.07 | 1.00 ± 0.45 | 285.97 ± 84.70 | 10 |
| 6397 | DL | 3.54 ± 3.20 | 0.61 ± 0.26 | 0.11 ± 0.17 | 1.64 ± 1.12 | 153.96 ± 45.83 | 10 |
| 6319 | DL | 7.37 ± 2.06 | 0.36 ± 0.18 | 0.13 ± 0.07 | 1.18 ± 0.40 | 268.36 ± 95.41 | 15 |
| 6282 | AS | 5.63 ± 2.42 | 0.35 ± 0.19 | 0.17 ± 0.12 | 1.45 ± 0.69 | 325.90 ± 127.35 | 9 |
| 6315 | DL | 5.91 ± 2.91 | 0.55 ± 0.22 | 0.21 ± 0.17 | 1.27 ± 0.65 | 350.60 ± 89.84 | 11 |
| 6302 | AS | 7.91 ± 2.98 | 0.71 ± 0.18 | 0.24 ± 0.11 | 1.36 ± 0.50 | 299.0 ± 96.53 | 13 |
| 6318 | DL | 5.55 ± 3.14 | 0.62 ± 0.27 | 0.40 ± 0.18 | 2.00 ± 1.10 | 265.73 ± 85.21 | 14 |
| 6310 | DL | 6.55 ± 4.03 | 0.69 ± 0.22 | 0.40 ± 0.19 | 1.18 ± 0.60 | 347.72 ± 105.52 | 14 |

**Table 3.**
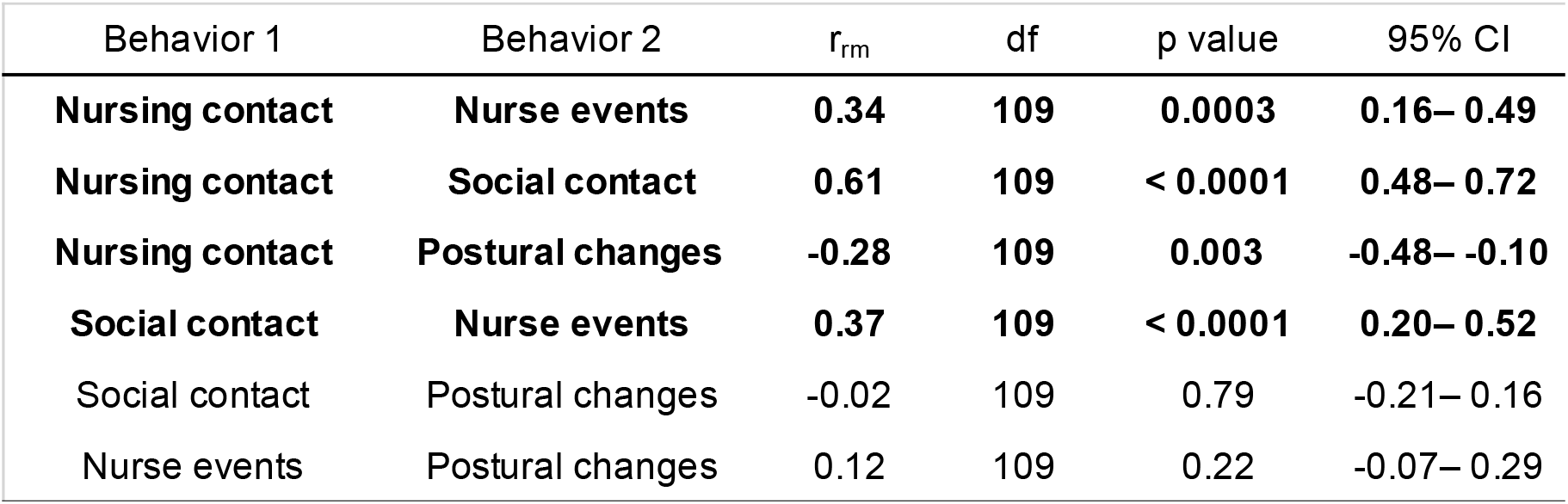
Repeated measures correlations between maternal behaviors. Bolded correlations are significant at the Bonferroni corrected threshold of p < 0.008.

| Behavior 1 | Behavior 2 | $r_{rm}$ | df | p value | 95% CI |
| --- | --- | --- | --- | --- | --- |
| **Nursing contact** | **Nurse events** | **0.34** | **109** | **0.0003** | **0.16– 0.49** |
| **Nursing contact** | **Social contact** | **0.61** | **109** | **< 0.0001** | **0.48– 0.72** |
| **Nursing contact** | **Postural changes** | **-0.28** | **109** | **0.003** | **-0.48– -0.10** |
| **Social contact** | **Nurse events** | **0.37** | **109** | **< 0.0001** | **0.20– 0.52** |
| Social contact | Postural changes | -0.02 | 109 | 0.79 | -0.21– 0.16 |
| Nurse events | Postural changes | 0.12 | 109 | 0.22 | -0.07– 0.29 |

### Salivary oxytocin and maternal care

We measured salivary OXT concentrations in 119 samples (10-11 samples per sow) following behavioral observations. Sow average OXT concentrations were 262.8 ± 107.2 pg/mL. The nursing contact model differed significantly from a reduced model that included only the control predictors (chi-square = 4.35, df = 1, p = 0.036). Results of the model indicate that nursing contact significantly predicted salivary OXT concentrations (est ± SE = 0.87 ± 0.35, p = 0.036, Fig 2). None of the control predictors influenced salivary OXT (Table 4). The second model considering all social contact between sows and piglets did not differ from the reduced model containing only control predictors (chi-square = 1.71, df = 1, p = 0.19). These results indicate that salivary OXT in these sows related to their extent of nursing contact specifically and not to more general social contact with piglets.

**Table 4.** Salivary oxytocin concentrations in sows are influenced by their nursing contact. Results of a LMM including control predictors. Significant predictors are indicated in bold. Sqrt= square root transformed

| Predictor | Estimate ± SE | chi-square | df | p value |
| --- | --- | --- | --- | --- |
| (Intercept) | 15.87 ± 0.55 | | | |
| **sqrt(proportion nursing contact)** | **0.87 ± 0.35** | **4.35** | **1** | **0.037** |
| sqrt(time delay nursing to sample) | -0.01 ± 0.26 | 0.00 | 1 | 0.978 |
| time of sample collection | 0.34 ± 0.25 | 1.61 | 1 | 0.204 |
| days since birth | 0.20 ± 0.33 | 0.31 | 1 | 0.579 |

**Fig 2.**
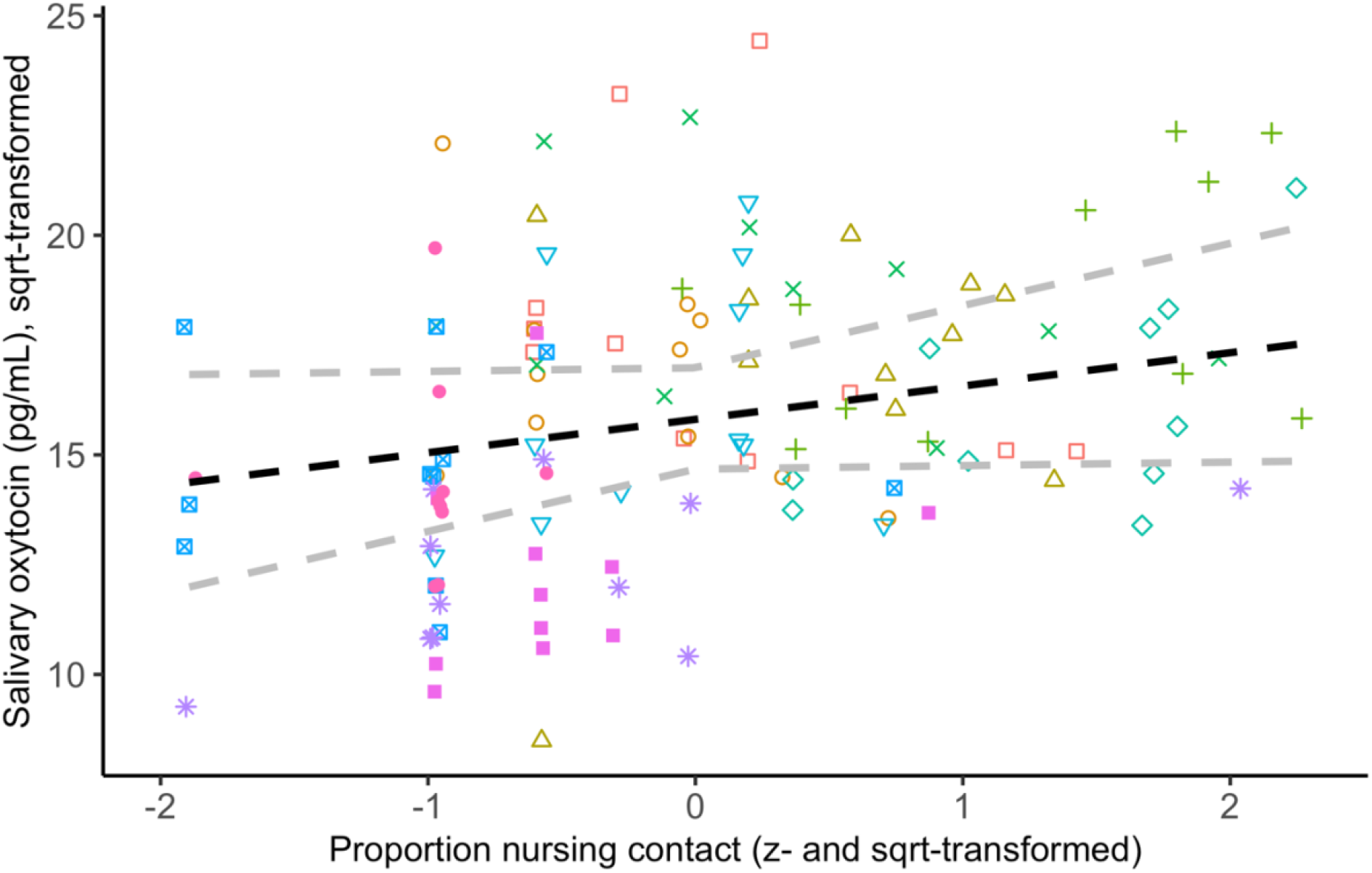
Relationship between salivary oxytocin and nursing contact. Sow identities are indicated by different shapes and colors. The dashed line indicates the expected values and 95% confidence intervals based on the fitted model (see Table 4).

## Discussion

Characterizing behavioral and physiological correlates of maternal care in domestic pigs can help to identify candidate traits that could be included in breeding programs to improve the welfare of sows and their piglets [9]. In one study that demonstrates this point, when a pig breeding line was established based on high post-natal survival rates, the high performing sows exhibited variation in several maternal behaviors compared to average performing sows in as quickly as two generations [50]. While some of these behavioral differences appeared to support offspring welfare, for example by reducing the likelihood of crushing behavior, high performing sows in indoor loose farrowing systems also exhibited *more* aggressive behavior towards their offspring, including savaging of offspring, compared to sows with average post-natal survival rates [51]. It would be interesting to explore whether selection criteria emphasizing behavioral and physiological indicators of maternal care in addition to indicators of piglet survival may result in better welfare outcomes for mothers and their offspring.

Behavioral data can be time-consuming to collect in livestock production settings [52], and complex behaviors such as maternal care are sometimes scored subjectively based on caretaker assessments [9,53]. Group scan sampling offers a fast and efficient method to measure behavior in multiple subjects within a short time-frame, and has been useful in identifying biologically-meaningful variation in maternal care in large rodent colonies [54]. A similar approach could be applied in conventional farrowing settings to better identify behaviors associated with different mothering styles in pigs. The sows in our study exhibited a great degree of inter-individual variation in their amount of nursing contact with piglets, which was stable within subjects during the later lactational period. Hourly nursing contact was positively correlated with the number of nursing events (when most piglets were in brief nursing contact), which are likely to indicate milk let-down [12]. Nursing contact was negatively correlated with the frequency of postural changes, which have been associated with greater risks of piglet crushing in other studies [55,56]. We consider nursing contact to require tolerance from sows, since nursing behavior is under their control, and sows can easily block access to their teats either by standing up (when piglets are smaller) or by laying on their stomachs. It is likely that prolonged contact with teats outside of brief milk-letdown events serves multiple functions, including potentially priming future milk-letdown events [15], as well as facilitating social bonding [57]. Our results suggest that the extent of nursing contact during lactation may be a good indicator of maternal style and maternal investment more generally in pigs.

Identifying physiological correlates of maternal behavior in sows can help to understand proximate causes of variation in maternal investment and may provide a complementary route to identify selection criteria for enhanced maternal care. In one of the few studies to investigate OXT related to mother-offspring social interactions in pigs, Valros and colleagues [22] did not find any relationship between plasma OXT concentrations in sows and their frequencies of affiliative social interactions with offspring. However, behavioral data and plasma samples were taken only once per subject, and on different days, which may have interfered with the ability to detect such links if they exist.

Salivary OXT measurements offer a minimally-invasive alternative to plasma that is amenable to repeated sampling while reducing interference with on-going behavior. Some researchers have even suggested that saliva may represent a superior matrix for measuring peripheral OXT, since blood and plasma contain proteins that may interfere with detection, and urine has a high salt concentration that may interfere with measurement [35,58]. However, further validations are needed to determine best method practices for treatment of saliva samples prior to measurement, especially regarding the need for extractions [36,37], which may depend on the species of interest [36]. More research is also needed to determine exactly what portion of OXT is measured in saliva (e.g. bound vs. free, intact vs. metabolites), which may depend on the specific assay and antibodies used [35,38]. We agree with Maclean and colleagues [35] that there may be more than one appropriate method for measuring salivary OXT, but this only increases the importance of reporting full analytical validations of new approaches when applied to new species [59,60]. In addition, more biological validations that measure salivary OXT related to events that are mediated by the oxytocinergic system (e.g. mating, parturition, lactation) or following pharmacological manipulations (e.g. administration of OXT agonists or antagonists), are greatly needed. Reporting the results of such studies, including those that find meaningful co-variation in endogenous OXT measurements and those that do not, is important to further validate measurements of salivary OXT, and to determine the extent and limits of its usefulness as a non-invasive biomarker of OXT-system activation.

Here we validated a method to measure salivary OXT in pigs that supports some previous results. Similar to Lürzel and colleagues [36], but in contrast with other studies [33,37], we found low correlations between extracted and unextracted saliva samples in pigs and believe that extraction may have reduced potential interference from matrix effects, consistent with evidence in plasma samples [24,59]. Similar to MacLean and colleagues [33], we found that the Caymen assay performed better than other commercially available assays for measuring salivary OXT. While measurements of duplicate saliva samples in Arbor and Caymen assays were generally in agreement, parallelism was less accurate, and reproducibility was lower in the Arbor assay. It should be noted that the extractions for both assays were conducted using the same SPE protocol, and as a result we did not use the extraction solution provided by the Arbor assay. While SPE is also a suitable option for use with the Arbor assay, it is possible that the use of their assay-specific extraction solution may improve performance of that assay.

Our results indicate that salivary OXT co-varies with parturition and nursing behavior and may be useful as an indicator of differences in maternal investment in sows. Salivary OXT increased nearly three-fold during parturition, although there was also a great deal of inter-individual variation in OXT responses, which is consistent with variation in plasma OXT responses to parturition in sows [17]. This variation may be explained in part by individual differences in baseline oxytocin concentrations as well as the duration of parturition, or by differences in the timing of saliva sampling relative to the timing of OXT-induced contractions. More research is needed to determine the extent and causes of inter-individual variation in peripheral oxytocin concentrations in different matrices, under baseline conditions and in response to biologically-meaningful events.

Sow salivary OXT concentrations during lactation co-varied with their extent of nursing contact with piglets. Although nursing contact was positively correlated with our measure of total social contact between sows and piglets, we did not find any relationship between salivary OXT and total social contact. It thus remains unclear whether salivary OXT is a useful indicator of mother-offspring social interactions more generally, outside of nursing contexts, in sows. In the only other study to measure salivary OXT in sows at multiple time points during lactation [37], sows varied greatly in salivary OXT concentrations on post-partum day 1, but their maternal behavior was not measured, and sows were also administered OXT i.m. on post-partum day 1, so it seems likely that salivary OXT measurements on that day were related to exogenous OXT administration, rather than to the sows’ behavior.

While sows had full control over their extent of nursing contact with piglets, the design of the conventional farrowing pen reduced their control over more general social contact with piglets. Early post-partum, while their movement was partially restricted by a restraining bar, sows had limited ability to initiate or avoid contact with piglets. Once the restraining bar was removed, sows could move freely in the pen, but could not initiate contact with piglets when they were in their laying area. As a result, throughout the lactational period piglets had more control over their extent of physical contact with sows than vice versa. It would be interesting to follow-up by recording similar behavioral and physiological measures of maternal care in alternative farrowing systems [61] in which several lactating sows are sometimes housed together in larger stalls where they have more control over their social contact with other sows and with piglets [62,63]. It is possible that salivary OXT may co-vary with more general mother-offspring social contact in settings where sows have more control over their social interactions.

The high frequency of mother-offspring interactions in the conventional farrowing pen also made it difficult to collect suitable baseline or pre-nursing samples without separating sows from their offspring, which would have interfered with the maternal behavior that we wanted to study. We are currently conducting further studies in weaned pigs in which we can more readily compare changes in salivary OXT related to a broader range of positive, negative and neutral social contexts. This next step will help to further investigate the usefulness of salivary OXT as a more general indicator of social affiliation and affective states in pigs.

## Supporting information

Moscovice etal_Salivary OXT Supplemental files

## Acknowledgements

We are grateful to Marianne Zenk for her help with the coordination of this project and for providing relevant information on pig husbandry. We would like to thank Dirk Ameling, Herbert Asmus, Frank Hintze and Heidi Schumann for their daily care of the pigs at the Experimental Pig Facility. We gratefully acknowledge technical support from Corinna Bochat with sample collection, and from Frieder Hadlich with data visualization.

