## Supplementary material for "Salivary oxytocin co-varies with parturition and nursing behavior in domestic pigs (*Sus scrofa domesticus*)": Moscovice etal_Salivary OXT Supplemental files

Supporting Information

**S1 Table**. **Stability in maternal behaviors during lactation.** Bolded Spearman correlations are significant at the Bonferroni corrected threshold of p < 0.004. Time 1= days 1-10 post-birth, Time 2= days 11-19 post-birth, Time 3= days 20-27 post-birth.

| Lactational period | Maternal Behavior | r_s_ | p value |
| --- | --- | --- | --- |
| Time 1 vs. Time 2 | nursing events | 0.46 | 0.15 |
| Time 1 vs. Time 3 | nursing events | 0.31 | 0.35 |
| Time 2 vs. Time 3 | nursing events | 0.33 | 0.32 |
| **Time 1 vs. Time 2** | **total contact** | **0.91** | **< 0.0001** |
| Time 1 vs. Time 3 | total contact | 0.70 | 0.02 |
| Time 2 vs. Time 3 | total contact | 0.68 | 0.03 |
| Time 1 vs. Time 2 | postural changes | 0.08 | 0.83 |
| Time 1 vs. Time 3 | postural changes | 0.05 | 0.90 |
| Time 2 vs. Time 3 | postural changes | 0.77 | 0.01 |
| Time 1 vs. Time 2 | nursing contact | 0.68 | 0.03 |
| Time 1 vs. Time 3 | nursing contact | 0.64 | 0.04 |
| **Time 2 vs. Time 3** | **nursing contact** | **0.97** | **< 0.0001** |

**S1 Fig.** **Comparison of parallelism obtained in EIAs from Caymen Chemicals (a) and Arbor Assays (b).** Red circles indicate the standard curve using the kit OXT standard, and blue triangles indicate different dilutions of extracted pooled saliva samples. b/bo= sample optical density / maximum binding optical density.

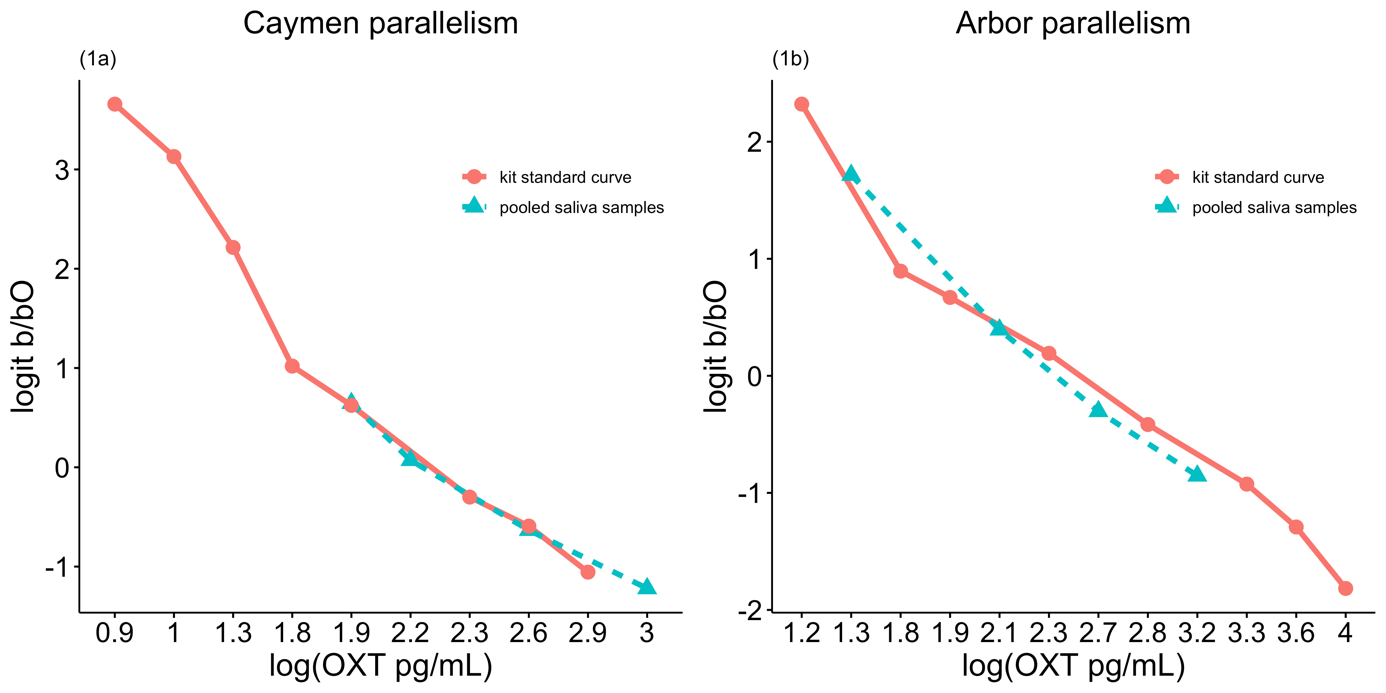
